# Immunofluorescence staining protocol for STED nanoscopy of *Plasmodium*-infected red blood cells

**DOI:** 10.1101/416776

**Authors:** Ann-Kathrin Mehnert, Caroline Sophie Simon, Julien Guizetti

**Affiliations:** Centre for Infectious Diseases, Parasitology, Heidelberg University Hospital, Im Neuenheimer Feld 324, 69120 Heidelberg, Germany

**Keywords:** Malaria, *Plasmodium falciparum*, Blood stage, Immunofluorescence, STED, Super-resolution

## Abstract

Immunofluorescence staining is the key technique for visualizing organization of endogenous cellular structures in single cells. Labeling and imaging of blood stage *Plasmodium falciparum* has always been challenging since it is a small intracellular parasite. The gold standard for parasite immunofluorescence is fixation in suspension with addition of minute amounts of glutaraldehyde to the paraformaldehyde-based solution. While this maintains red blood cell integrity, it has been postulated that antigenicity of the parasite proteins was, if at all, only slightly reduced. Here we show the deleterious effect that even these small quantities of glutaraldehyde can have on immunofluorescence staining quality and present an alternative cell seeding protocol that allows fixation with only paraformaldehyde. The highly improved signal intensity and staining efficiency enabled us to carry out RescueSTED nanoscopy on microtubules and nuclear pores and describe their organization in greater detail throughout the blood stage cycle.

**Highlights:** - Omitting glutaraldehyde from fixative improves immunofluorescence staining

- STED nanoscopy is readily applicable to infected red blood cells

- Intranuclear microtubules are nucleated from distinct sites

---

Over the last decade significant progress in understanding the cell biology of malaria parasites has been made [1]. Despite advances in live cell tagging technologies, immunofluorescence staining is still the best alternative to localize unaltered endogenous proteins in their native context [2]. The red blood cell (RBC) stage of the *Plasmodium falciparum* parasite is particularly challenging to stain due to multiple membranes hindering antibody penetration, the autofluorescence of the host cell, and its sensitivity to osmotic changes, causing lysis under many conditions. Tonkin *et al.* have made a great effort in determining the minimally required glutaraldehyde (GA) concentration (0.0075%) to prevent RBC lysis throughout the whole in suspension immunofluorescence staining procedure [3]. Paraformaldehyde (PFA) fixation alone is not a viable option for immunofluorescence staining in suspension because RBC lysis after or during permeabilization makes the recovery of parasites impossible. This protocol is widely used in the field and has significantly contributed to the progress around many diverse cell biological questions. However, GA fixation remains a compromise. Its deleterious effects on immune epitopes, quenching of fluorescent proteins, and increase of background fluorescence at higher concentrations are well-known. It has for example been reported that even minute amounts of GA prevent recognition of the plasmodial Rex2 [4]. Alternative fixation methods such as methanol, air drying, or saponin-lysed parasites have been used but are clearly inferior in terms of structural preservation. Also glyoxal, which has been recently highlighted as a superior fixative for many vertebrate and mammalian cell lines in terms of structural preservation and staining efficiency [5], did not work efficiently in *Plasmodium* blood stage (Suppl. Fig. 1 & Suppl. Table 1). Another key limitation to studying parasite cell biology is its comparatively small size. There might, however, be intriguing structural features such as microtubule reorganization throughout schizogony and nuclear pore distribution, which are below the diffraction limit [6]. This limitation has been partially overcome by applying structured-illumination microscopy (SIM) during invasion studies [7]. SIM is however physically limited to a doubling of resolution [8]. More advanced super-resolution imaging technologies, such as Stimulated Emission Depletion (STED) heavily rely on optimal sample preparation, good signal-to-noise ratio, and high signal intensities [9,10]. Unfortunately high STED depletion laser intensities cause cell destruction when infected red blood cells with hemozoin are imaged and so far only hemozoin free stages such as liver cells and merozoites have been imaged using STED [11,12]. Here, we describe the effect of omitting GA from the fixation protocol by combining immunofluorescence with cell seeding on imaging supports as described previously [13–16]. This resulted in a substantial improvement of labeling intensity, specificity and efficiency while maintaining structural integrity of the parasite cells. Additionally, we show that antibodies that have been discarded as unsuitable for *P. falciparum* staining can actually produce satisfactory results under those conditions. The described immunofluorescence and imaging protocol enabled us also to acquire high quality STED pictures of entire iRBCs showing individual nuclear pore clusters and microtubule reorganization.

To avoid issues with RBC lysis in suspension we used the possibility to seed the cells in concanavalin A-coated imaging dishes, as described by Grüring *et al.*, prior to fixation and staining [16]. This prevented parasite loss during fixation with 4% PFA only and allowed us to explore the effect of 0.0075% GA addition on the quality of immunofluorescence staining. Without GA we observed a change of the infected RBC (iRBC) monolayer from red-brown to clear after the permeabilization step, suggesting that the RBC cytoplasm was at least partially washed out. We found that RBCs were structurally slightly altered and became significantly more translucent in the brightfield channel (Fig. 1A). Host cell actin staining with phalloidin revealed a more irregular RBC membrane shape whereas the staining intensity was much increased. However, we did not observe any loss of parasite density on the slide. From a practical aspect, this staining technique achieves very consistent monolayers with high parasitemia, avoids centrifugation steps, and facilitates potential combination with live cell imaging. Using parasitophorous vacuole staining with anti-Exp1 antibodies [17], we observed that the three-dimensional structure of the parasite showed neither structural alterations nor flattening of the parasite itself or its nucleus (Fig. 1B). This is consistent with the observation that partial RBC lysis only occurred well after fixation was completed, allowing the parasite to maintain its shape. While this could limit detection of RBC cytoplasm proteins, we observed, however, strongly increased Exp1 signal intensity without GA and a much higher labeling density. When the same cell was z-projected, the anti-Exp1 signal is only barely detectable in the GA-fixed samples under identical acquisition and contrast settings (Fig. 1C). This prompted us to test this improved staining protocol for several antibodies targeting different parasitic structures representing the Maurer’s clefts, the parasite outer membrane, parasite cytoplasm, the nuclear membrane and the nucleoplasm. Labeling with anti-α-tubulin antibodies revealed much brighter staining of nuclear microtubules under identical conditions while displaying the same protein organization (Fig. 1D). When fixed in the presence of GA, both centriolar plaques and hemi-spindle microtubules were only detectable if contrast settings for the anti-α-tubulin signal were adjusted accordingly (Fig. 1D). Anti-Nup116 (nuclear pores, [18]) staining in contrast yielded comparable signal intensities when fixed either with or without GA (Fig. 1E). Staining with anti-SBP1 (Maurer’s clefts, [19]) antibodies, although the same localization pattern was maintained, was much more efficient without GA in all cells observed (Fig. 1F). Our results suggest that even those small quantities of GA can cause a significant decline in antibody binding.

**Figure 1.**
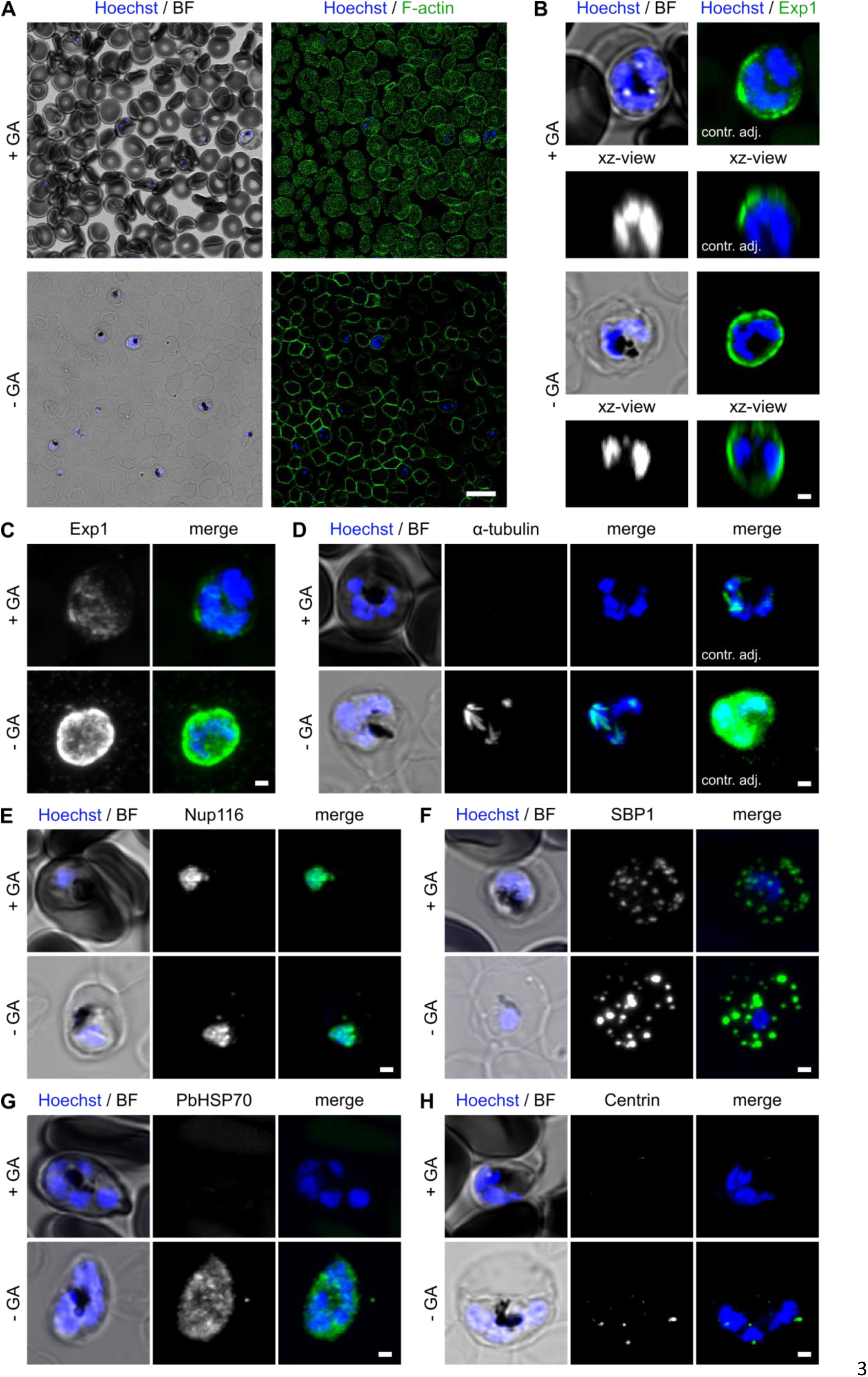
Infected RBC morphology and immunofluorescence staining efficiency after fixation with or without GA. (A) Confocal overview images of iRBCs stained with Hoechst and Phalloidin-Alexa Fluor 488 for F-actin after fixation with or without addition of GA. Scale bar, 10 µm. (B) Confocal slice images of iRBCs stained with Hoechst and labeled with anti-Exp1 antibody. To illustrate the three-dimensional shape of the parasite, xz-views of the same cells are given in each panel. Contrast adjustments are different for +GA and – GA samples to allow visibility of the signal. (C)–(H) Confocal images of iRBCs, fixed either with or without GA addition, stained with Hoechst and labeled with antibodies for various parasitic structures. All images except brightfield (BF) are maximum intensity projections. Identical acquisition and contrast settings were applied, except where indicated (contr. adj.). (C) Exp1 (parasitophorous vacuole). Z-projection of the same cell as in (B) without the previous contrast adjustments. (D) α-tubulin (microtubules). (E) Nup116 (nuclear pores). (F) SBP1 (Maurer’s clefts). (G) PbHSP70 (parasite cytoplasm). (H) Centrin (centriolar plaques). Scale bars (B)–(H), 1 µm.

More critically, we detected, as previously found by many colleagues, a high fraction of unlabeled cells (Suppl. Fig. 2). Quantification revealed a strong drop of labeling density with GA for PbHsp70, Exp1, SBP1, and Cdc48 (apicoplast, [20]) antibodies (Table 1) even though we used standard incubation conditions and Triton^®^ X-100 concentration as described before [3]. We, however, noted a strong enrichment of double- and triple-infected RBCs in the stained fraction of +GA-treated cells. To exclude the possibility that the lack of staining in GA-fixed cells is related to the absence of mechanical stress imposed upon cells during centrifugation, potentially improving antibody penetration, we also tested staining in suspension as described previously [3]. This approach did not improve the fraction of stained cells (Table 1). A possible explanation for the increase in labeling density under GA-free conditions may be reduced epitope masking or improved penetration of the antibody to the site of binding due to partial lysis of RBCs. To distinguish between those possibilities we pretreated seeded cells with Saponin before fixation and then stained for SBP1. This rescued the drop of labeling density, whereas treating the cells with Saponin after fixation, did not (Suppl. Fig. 3). We conclude that penetration is the main limiting factor for labeling density. This poses the question whether an unbiased selection of cells for quantification purposes is at all possible using fixation protocols with GA.

**Table 1.** Percentage of antibody-stained cells with or without GA addition. Confocal overview images of iRBCs stained with Hoechst and respective antibody after fixation with or without addition of GA in Lab-Teks and with GA in suspension were quantified for presence or absence of antibody staining after applying uniform contrast threshold. Number of positive cells over total counted cells in brackets.

| Fixation\Antibody | anti-PbHSP70 | anti-Exp1 | anti-SBP1 | anti-Cdc48 |
| --- | --- | --- | --- | --- |
| -GA on Lab-Tek | 96.4% (187/194) | 99.5% (204/205) | 99.5% (189/190) | 100% (132/132) |
| +GA on Lab-Tek | 0% (0/92) | 4.7% (9/191) | 2.6% (3/117) | 8.5% (5/59) |
| +GA in suspension | 0% (0/62) | 2.7% (2/75) | 2.7% (2/73) | 3.2% (2/62) |

To check whether GA can occasionally prevent antibody recognition, we tested an anti-PbHSP70 and anti-Centrin antibody that had been previously used to stain methanol but not PFA+GA fixed *P. falciparum* blood stage parasites [21,22] We confirmed that anti-PbHSP70 did not label cells with GA (Table 1) but could give a distinct cytoplasmic staining without (Fig. 1G). Staining with commercial cross-species reactive anti-Centrin antibody revealed only background staining with GA (Fig. 1H). Under improved conditions we detected clear perinuclear localization at a site that is presumably the centriolar plaque, thereby displaying the expected localization.

The high signal-to-noise ratio achieved by the described staining protocol is particularly beneficial for reaching improved image quality in super-resolution microscopy [9]. We therefore imaged our samples by STED. An important limitation upon parasite imaging is substantial structural damage to the cells caused by the high-power STED laser should it hit the highly absorbing hemozoin crystal [23]. This effect has been described before by Michael Pasternak (https://www.abberior-instruments.com/products/expert-line/rescue/). Hence we implemented an adaptive illumination module named RescueSTED, which automatically, based on a threshold intensity setting, inactivates the STED laser in low fluorescence regions, comprising the hemozoin. This allowed us to resolve sub-diffraction organization of microtubules within entire infected RBCs of asexual blood stages (Fig. 2A). While rings had no detectable tubulin structure a small fraction of trophozoites displayed either perinuclear tubulin clusters or intranuclear microtubule structures, termed hemispindles (Fig. 2B). This suggests they are close to the first mitotic division. Interestingly, hemispindle microtubules sometimes extend way beyond the boundary of nuclear staining, which is frequently accompanied by protrusions in the Hoechst (data not shown). STED images, in contrast to confocal, clearly show that hemispindle microtubules are nucleated at distinct sites just below a groove in Hoechst staining (Fig. 2A). This points towards a centriolar plaque organization that is clearly distinct from the microtubule organizing centers found in e.g. yeast or other model organisms. These hemispindles and tubulin clusters were more frequently found in schizont nuclei showing that they are intermediates of microtubule reorganization (Fig 2A-B). In 52% of schizonts both of these intermediates were present in different nuclei, highlighting the asynchronous nature of nuclear division in *P. falciparum* [24]. In segmenting schizonts, after nuclear division is completed, STED microscopy revealed the bilayered structure (<100nm distance) of the microtubule network underlying the inner membrane complex of separating merozoites. To more clearly demonstrate the association of schizont microtubules with the nucleus we outlined the parasite cytoplasm by labeling the parasitophorous vacuole protein Exp1 (Fig. 2C). This also shows the feasibility of dual-color STED although the ATTO 647 labeled secondary antibody performed less well than the ATTO 594 labeled one. To further validate our approach we attempted to image individual nuclear pore clusters using anti-Nup116 staining in early- and late-stage mononucleated parasites (Fig. 2D). Despite the Nup116 antibody not being entirely devoid of unspecific background staining and a likely underestimation of nuclear pore number due to potentially close clustering, the amount of counted nuclear pore clusters matched well with the numbers previously described in an electron microscopy study (Fig. 2E) [18,25].

**Figure 2.**
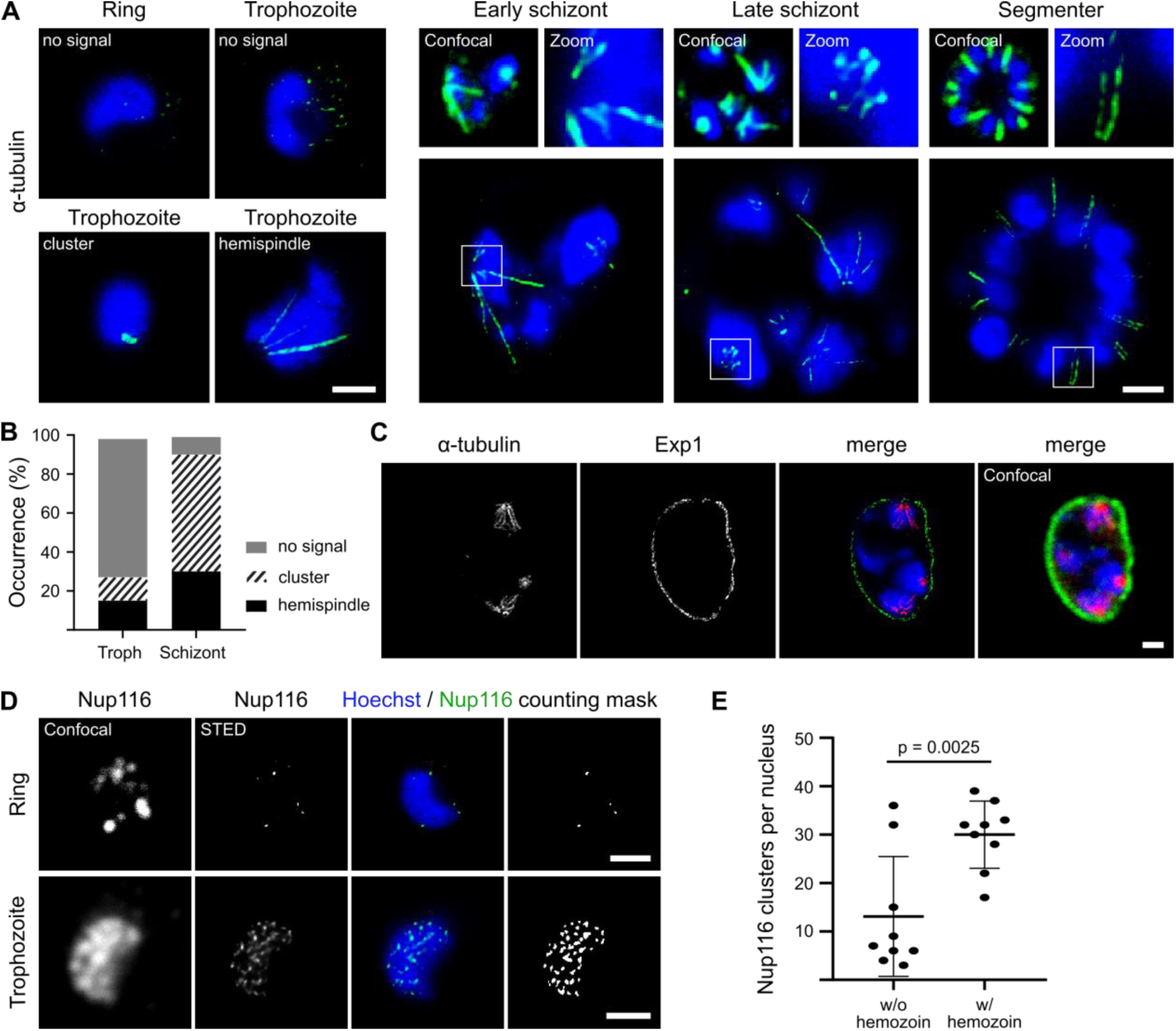
STED nanoscopy images of parasite nuclear structures. (A) Images acquired by RescueSTED showing parasite nuclei and observed tubulin structures in different life cycle stages. Infected RBCs were stained with Hoechst and anti-α-tubulin antibody. For schizont stage confocal images and zoomed in regions are shown (B) Quantification of occurrence of nuclear tubulin structures if present, i.e. clusters and hemispindles by confocal microscopy. 131 trophozoites and 46 schizonts (129 nuclei) were analyzed for this purpose. (C) iRBCs were stained with Hoechst and labeled with antibodies for α-tubulin and Exp1 to highlight parasite boundaries and demonstrate intranuclear tubulin staining. Images are single slices acquired by RescueSTED. (D) RescueSTED images of iRBCs stained for nuclear pores by anti-Nup116 antibody. Nuclear pore clusters were quantified by automated segmentation (see counting mask panel) in mononucleated parasites without hemozoin, i.e. rings, and with hemozoin, i.e. trophozoites. Quantification results of nine representatives each are given in (E). Statistical significance analyzed by Student’s t-test. Scale bars (A), (C)–(D), 1 µm.

The amount and quality of cell biological information that can be extracted from fixed cells is highly dependent on the quality of the immunofluorescence staining. Our study demonstrates that fixation without GA improves specificity and efficiency of staining. An optimal staining protocol forms the basis of efficient super-resolution nanoscopy and enabled us to uncover previously unseen organizational details of microtubules and nuclear pores in malaria parasite blood-stages.

## Materials and methods

### Parasite culture

*P. falciparum* blood stage cultures (NF54) were maintained at a hematocrit of about 3% in 37 °C incubators with 90% humidity, 5% O_2_ and 3% CO_2_ saturation. Culture development was monitored by Giemsa staining, and cultures were diluted with O+ human erythrocytes to maintain a parasitemia of 3–5%. Parasites were synchronized by sorbitol lysis of late stages.

### Seeding infected red blood cells

Cell seeding was adapted to Lab-Tek II 8 well chambered slides (Thermo Fisher) using previously described procedure [16]. Infected RBCs containing *P. falciparum* ring stages were seeded on Lab-Tek II slides. For this purpose, Lab-Tek wells were coated with 80 µL Concanavalin A (Sigma, 5mg/ml in water) per well. After 20 min at 37 °C, wells were rinsed twice with pre-warmed PBS. For each well, 150 µL of iRBC culture was washed twice with PBS by centrifugation in 1.5 ml reaction tubes at 800 g for 15 sec and distributed into the wells. Cells were allowed to settle for 10 min at 37 °C. Unbound RBCs were carefully washed away with two to three PBS rinses until a faint RBC monolayer remained. Seeded iRBCs were incubated a few hours or overnight in 400µl complete RPMI cell culture medium, which was necessary since fixation of cells directly after iRBC seeding never allowed us to detect any spindle microtubules in the parasites (Suppl. Fig. 4). This effect is independent of the fixation procedure and might be due to depolymerization of these delicate structures upon osmotic stress caused by PBS or change in temperature during cell handling.

### Immunofluorescence staining

Wells were quickly rinsed with PBS and immediately fixed with prewarmed 4% PFA or 4% PFA/0.0075% GA at 37 °C for 20 min. All remaining steps were performed at room temperature. After rinsing once with PBS, cells were permeabilized with 0.1% Triton^®^ X-100 in PBS for 15 min, followed by three PBS washes. Quenching of free aldehyde groups was done in ~0.1 mg/mL NaBH_4_ for 10 min. Cells were washed twice and blocked in 3% BSA/PBS for at least 30 min. Both primary and secondary antibody dilutions (see Suppl. Table 2 for details) were centrifuged at 14.800 rpm for 10 min at room temperature to remove aggregates. Primary antibodies were allowed to bind for 2 h in 3% BSA/PBS followed by three washes with 0.5% Tween^®^ 20/PBS for 5-10 min each on a horizontal shaker. Secondary antibodies and Hoechst were added in fresh blocking buffer and incubated for at least 30 min while protected from light. Cells were washed again with 0.5% Tween^®^ 20/PBS three times and one additional time in PBS on a horizontal shaker. Samples were stored in 300 µL PBS at 4 °C in the dark until imaging. The same protocol was also applied to cells in suspension with 4% PFA/0.0075% GA fixation as described before [3]. RBC actin was stained with Phalloidin-Alexa Fluor 488 1:500 (Thermo Fisher) in PBS for 30 min. For antibody penetration test, cells were pre-treated with 0.05% Saponin for 30 sec at RT and then washed 0nce with PBS before fixation. Saponin post-treatment was carried out after fixation with 0.1% Saponin for 5 min at RT.

### Image acquisition and analysis

Images were acquired on a Leica TCS SP8 scanning confocal microscope, equipped with a 63x oil objective (numerical aperture, 1.4), GaAsP hybrid detectors and spectral emission filter. Laser power (405, 488, 561, 633) was individually adjusted to the different antibody combinations and fluorophores used. Images had a size of 128 x 128 pixels with a pixel size of 72 nm and pixel dwell time of 4.88 µs. Z-stacks of 4.18 µm and 6.27 µm were acquired with a step size of 280 nm. For overview images to quantify labeling efficiency we used 2048 x 2048 pixels with a pixel size of 92 nm and z-stack size of 8-10 µm. Images were merged and adjusted for contrast using Fiji [26]. Super-resolution images were acquired using a single-point scanning STED microscope by Abberior Instruments GmbH, equipped with a pulsed STED depletion laser of 775 nm, using gated detection. The STED laser power was adjusted to about 40%. Imaging was performed with a 100x oil objective (numerical aperture, 1.4) and a pixel size of 20 nm and pixel dwell time of 10 µs. To image entire iRBCs without cell destruction, we used the adaptive illumination module. RescueSTED was activated by CONF level, which defines the minimal intensity level to be reached in a confocal image before activating the STED laser at this pixel. CONF levels varied depending on sample and antibody between 20 to 100 and were adjusted if necessary for every image. Deconvolution was performed using Richardson-Lucy algorithm with default settings and regularization parameter of 10^-5^ in IMSpector imaging software (Abberior Instruments GmbH). Counting of Nup116 dots was done in Fiji using image segmentation (Auto Local Threshold method = Otsu, Radius = 5), Watershed, and Analyze Particles (size exclusion >4px²) functions. A maximum intensity threshold (>30) was applied to exclude segmented background staining.

## Acknowledgements

Funded by the German Research Foundation (DFG) – [349355339] to JG. We thank: Ann-Kristin Mueller for providing the anti-PbHSP70 and Jude Przyborski for the anti-SBP1, anti-Cdc48 and anti-Exp1 antibodies. Jude Przyborski and Markus Ganter for critical reading of the manuscript. Vibor Laketa, Head of the Infectious Diseases Imaging Platform (www.idip-heidelberg.org), for providing microscopy access and advice. Susann Kummer for advice on STED microscopy and sample preparation.

## Supplemental Figures and Tables

**Supplemental Figure 1.**
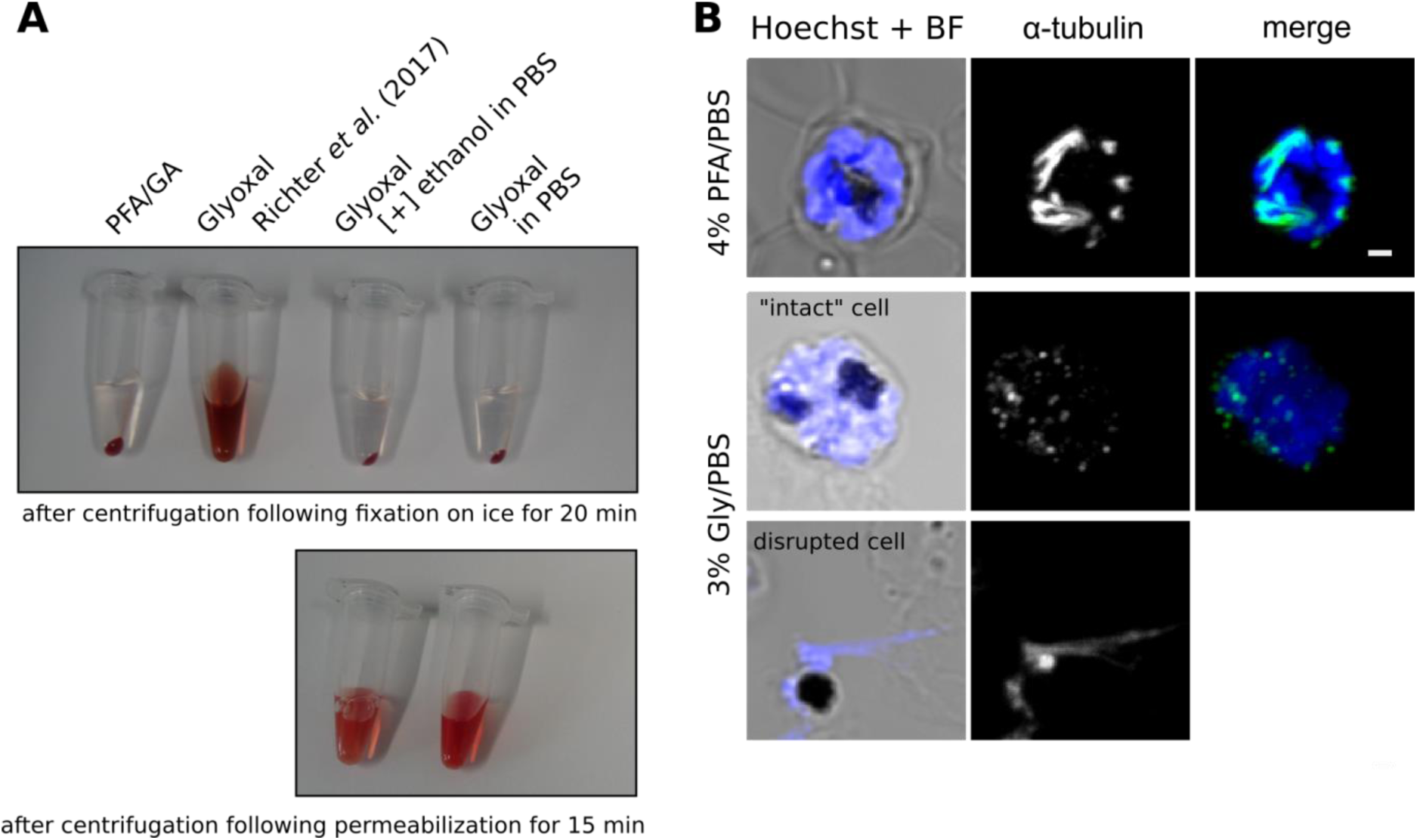
Fixation by glyoxal in solution and on Lab-Teks. Different glyoxal fixation solutions were tested on iRBCs in solution (A) or on Lab-Teks (B–C). If not indicated otherwise, fixation was performed on ice. (A) Exemplarily, the results of fixation by the following solutions are shown: 4% PFA/0.0075% GA in PBS, 3% glyoxal fixation solution according to Richter et al., 3% glyoxal/20% ethanol in PBS, and 3% glyoxal in PBS. Photos were taken after centrifugation following fixation on ice for 20 min (upper panel). Non-lysed suspensions were subjected to subsequent permeabilization and centrifugation (lower panel). (B) Confocal images of iRBCs fixed either by 3% glyoxal in PBS on ice or by 4% PFA in PBS on Lab-Teks at 37 °C. Cells were labeled with anti-α-tubulin antibody and DNA was stained with Hoechst. All images except brightfield (BF) are maximum intensity projections. Scale bar, 1 µm.

**Supplemental Table 1:**
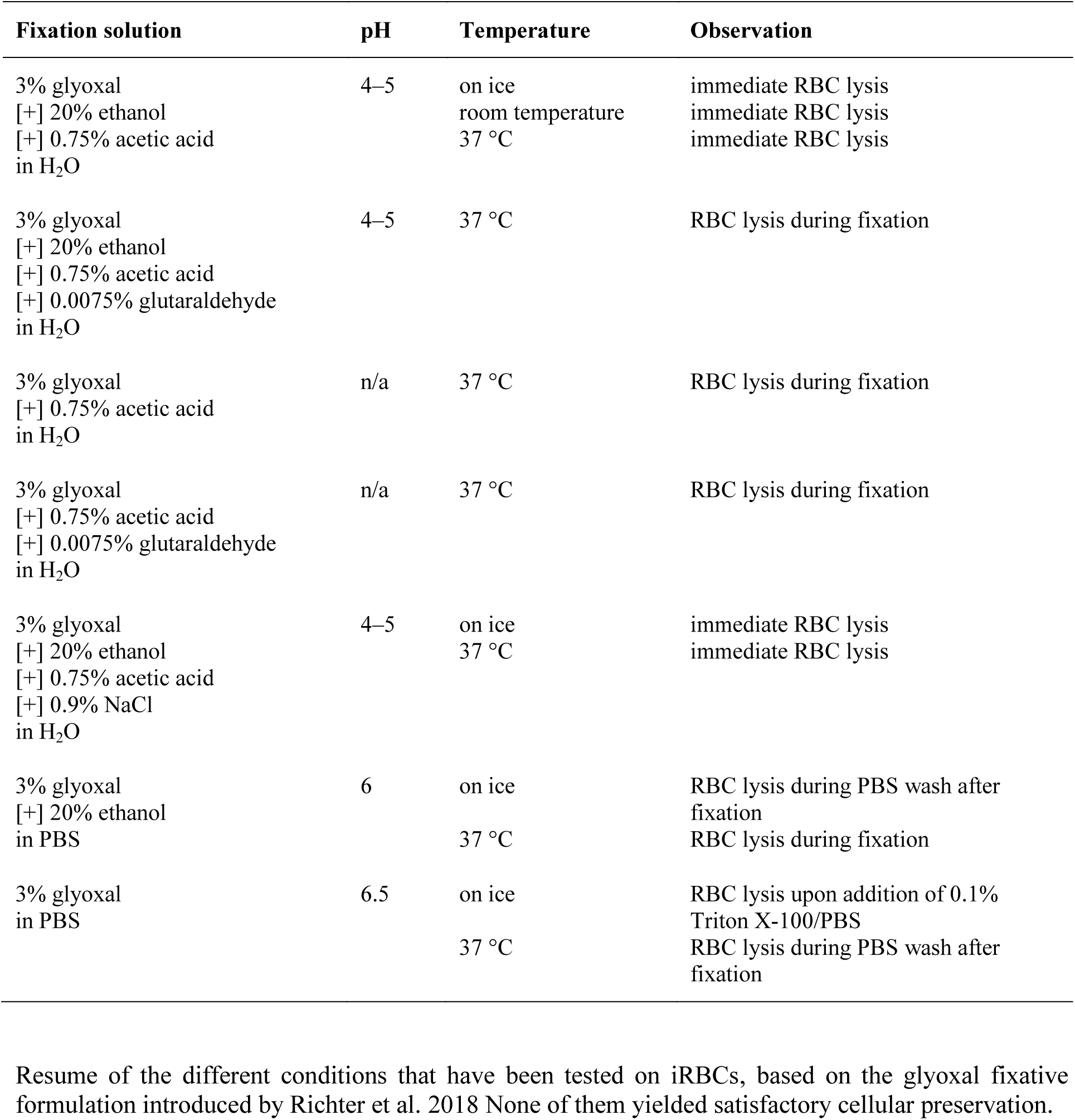
Tested glyoxal fixation solutions and conditions.

**Supplemental Figure 2.**
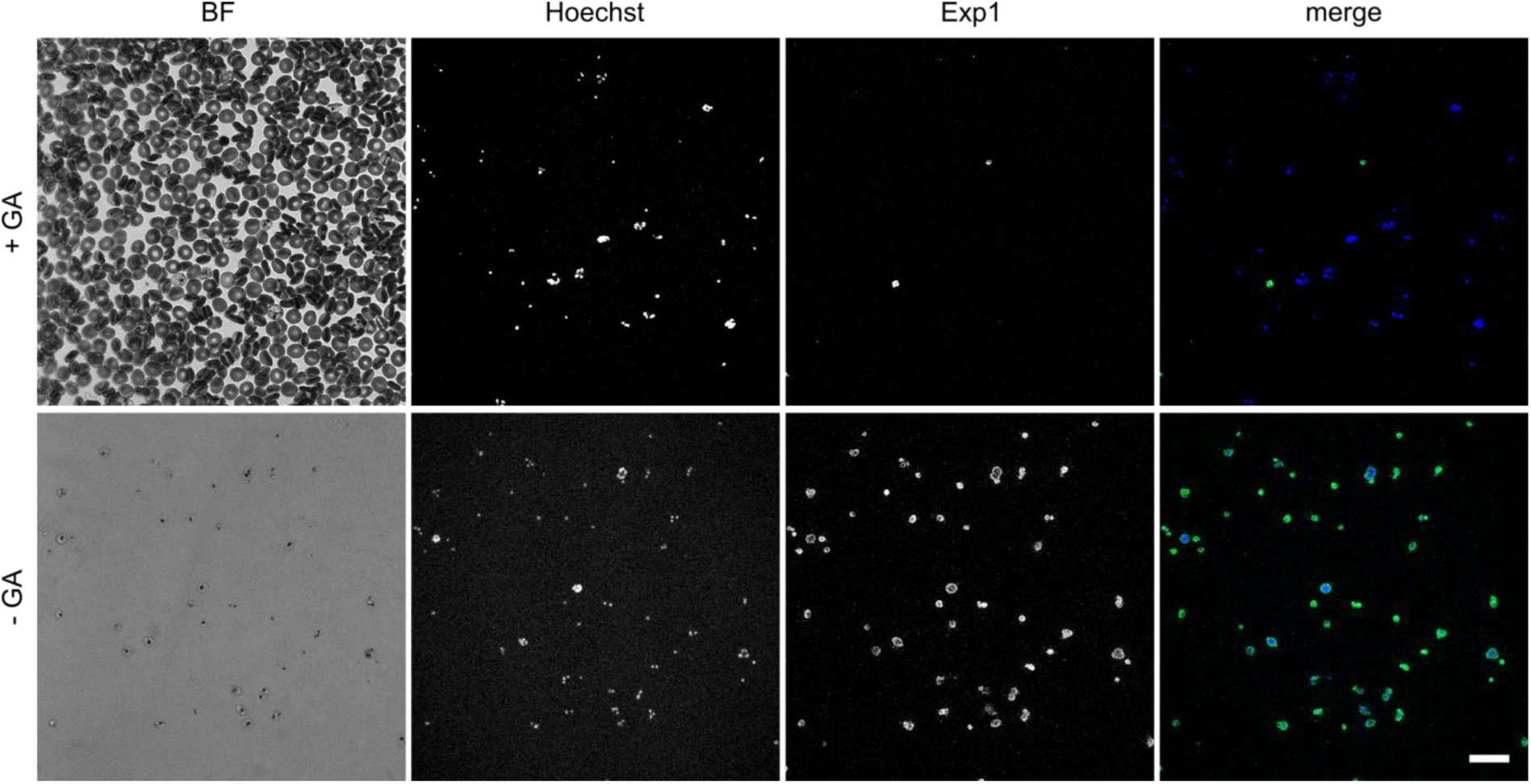
Labeling density of iRBCs with and without GA. Overview image of iRBCs stained with Hoechst (blue) and labeled with anti-Exp1 antibody (green) after either fixation with (upper panels) or without GA addition (lower panels). All images are maximum intensity projections and contrast adjustment is identical for both conditions. Scale bar, 20 μm

**Supplemental Figure 3.**
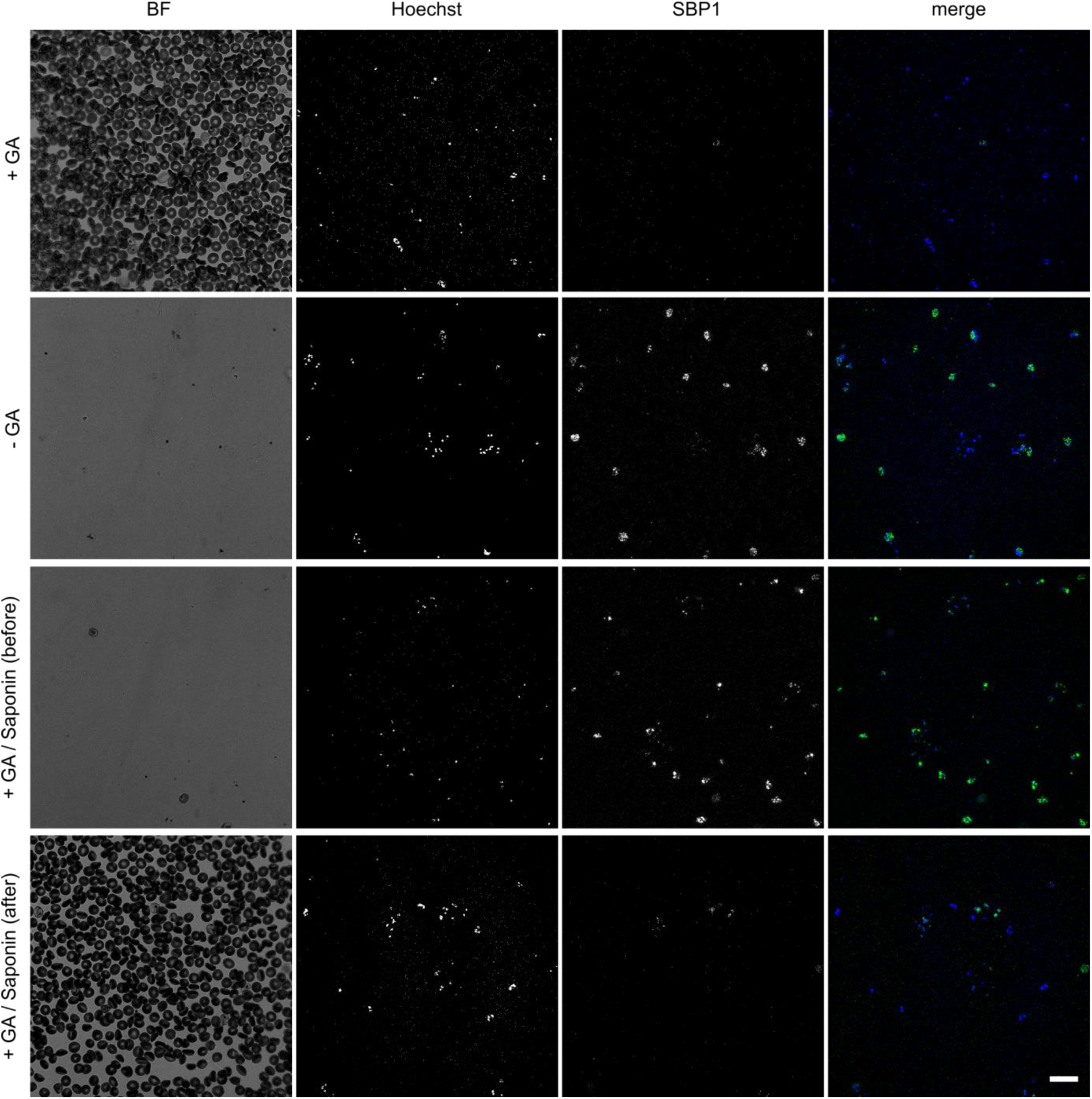
Labeling density of iRBCs with Saponin permeabilization. Overview image of iRBCs stained with Hoechst (blue) and labeled with anti-SBP1 antibody (green) after either fixation with GA, without GA, with GA pretreated with 0.05% Saponin 30 sec, and with GA posttreated with 0.05% saponin for 5min. All images are maximum intensity projections and contrast adjustment is identical for all conditions. Scale bar, 20 µm. Table below shows quantification of labeling density for all conditions.

| Fixation\Antibody | anti-SBP1 |
| --- | --- |
| -GA | 99.5% (189/190) |
| +GA | 2.6% (3/117) |
| +GA Saponin pretreatment | 89.8% (158/176) |
| +GA Saponin posttreatment | 7.8% (19/241) |

**Supplemental Figure 4.**
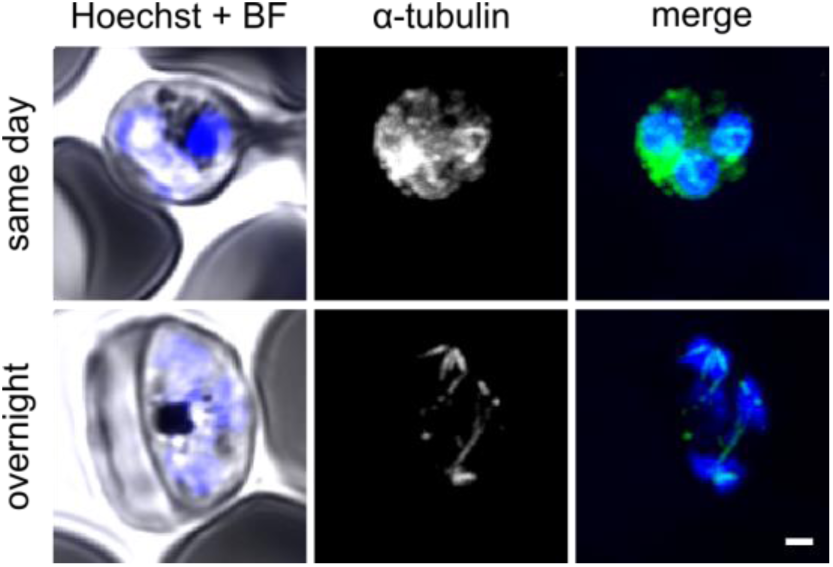
Overnight seeding is required to detect spindle microtubules. Infected RBCs were either seeded shortly before fixation (upper panel) or the day before (lower panel). Parasites were labeled with anti-α-tubulin antibody (green) to visualize mitotic spindles after fixation with GA. Image acquisition settings, i.e. laser power, were adjusted accordingly to allow signal visibility despite fixation with GA. All images except brightfield (BF) are maximum intensity projections. Scale bar, 1 µm.

**Supplemental Table 2:** List of used antibodies and dilutions.

| **Antibody target** | **Species** | **Source** | **Dilution** |
| --- | --- | --- | --- |
| Exp1 | Rabbit | Jude Przyborski | 1:500 |
| α-tubulin | Mouse | Sigma-Aldrich (T5168) | 1:500 |
| *PfNup116* | Rabbit | Julien Guizetti | 1:50 |
| SBP1 | Rabbit | Jude Przyborski | 1:500 |
| *PbHSP70* | Mouse | Ann-Kristin Mueller | 1:100 |
| Cdc48 | Rabbit | Jude Przyborski | 1:500 |
| Centrin | Mouse | clone 20H5, Merck (04-1624) | 1:1000 |
| Anti-mouse ATTO 594 | Goat | Sigma (76085-1ML-F) | 1:200–1:500 |
| Anti-rabbit ATTO 594 | Goat | Sigma (77671-1ML-F) | 1:200–1:500 |
| Anti-mouse ATTO 647N | Goat | Sigma (50185-1ML-F) | 1:200–1:500 |
| Anti-rabbit ATTO 647N | Goat | Sigma (40839-1ML-F) | 1:200–1:500 |

